# Select properties of supramammillary input to adult-born dentate gyrus neurons

**DOI:** 10.64898/2026.10.21.760824

**Authors:** Nirnath Sah, Olivia Curtis, Susan Lubejko, Alejandro Pinto, Shiliang Zhang, Marisela Morales, Carmen Vivar, Henriette van Praag

**Author notes:** Correspondence: Carmen Vivar, Henriette van Praag.

## Abstract

The hypothalamic supramammillary nucleus (SUM) projects to the hippocampal dentate gyrus (DG), a brain area crucial for learning, memory, and adult neurogenesis. How SUM inputs regulate adult-born DG (abDG) neurons to support memory processing remains unclear. Here, using monosynaptic tracing in adult mice, we identified a distinct subpopulation of abDG-targeting SUM neurons defined by large somata and low firing rate. Electrophysiological recordings revealed that these SUM neurons co-release glutamate and GABA onto abDGs, and that voluntary running selectively enhances GABAergic transmission. Chemogenetic silencing of SUM-DG GABAergic projections impaired pattern separation, the ability to make fine distinctions between similar stimuli, but not fear conditioning and spatial navigation memory. Altogether, our findings reveal that a distinct SUM subpopulation provides inhibitory modulation to abDG neurons, establishing a novel circuit mechanism for sparse encoding during pattern separation.

## Introduction

The hippocampus is essential for learning and memory (*1*). Within the hippocampus, the dentate gyrus (DG) gives rise to new neurons in adulthood (*2, 3*). Adult-born DG (abDG) neurons are highly regulatable by intrinsic and extrinsic factors; most notably, their number is strongly increased by exercise(*4*). The abDG neurons are considered to contribute to synaptic plasticity, spatial memory function (*5*), and pattern separation (*6, 7*), the ability to distinguish between highly similar events and stimuli (*8*). To better understand how new neurons modulate neuronal function, we and others have mapped their afferent network under sedentary and running conditions (*9–16*). Afferent inputs to abDGs originate primarily from the hippocampus and basal forebrain before expanding to include distal (sub)cortical areas, a pattern of connectivity that is approximately doubled by running (*16*). Unexpectedly, we observed that direct innervation of abDG neurons by the supramammillary nucleus (SUM) of the hypothalamus (*14*) is increased eight-fold by running wheel activity (*13, 16*). However, the properties of the SUM cells projecting to abDG neurons and the functional contribution of this input to behavior remain underexplored.

The hypothalamic SUM nucleus(*17*) consists of a heterogeneous neuronal population (*18–20*) implicated in social and spatial memory, sleep, addiction, and stress (*21*). Even so, the SUM contribution to episodic-like memory remains poorly understood(*22, 23*). It has been shown that SUM regulates the hippocampal theta rhythm (*24–29*). Activation of SUM-DG projections enhances theta synchronicity, whereas inactivation disrupts it (*28, 30*). Transient SUM silencing causes modest deficits in memory retention in the water maze and passive avoidance tests (*31–33*). More recent studies indicate that SUM projections to the DG modulate contextual novelty and novel place recognition memory (*34, 35*). In addition, SUM stimulation reportedly enhances neurogenesis and spatial memory, while reducing anxiety-like behavior (*36*). Neurotransmission from SUM onto the DG uniquely exhibits dual GABA and glutamate release (*19, 30, 34–40*). However, the specific circuit mechanism by which SUM input modulates abGCs to support hippocampal function remains poorly understood.

Here, using monosynaptic rabies virus tracing and patch-clamp recordings, we reveal that a specific subpopulation of SUM neurons forms functional connections onto abDG neurons co-releasing glutamate and GABA. Patch-clamp recordings following optogenetic stimulation of SUM axons revealed that voluntary running enhanced inhibitory transmission onto DG neurons. Furthermore, chemogenetic inhibition of SUM GABA release resulted in profound deficits in touchscreen pattern separation but not in spatial or fear memory performance. Altogether, these findings demonstrate that a specific SUM-DG circuit modulates abDG neurons, highlighting a prominent role for SUM GABAergic transmission in pattern separation, a cognitive process closely linked to adult neurogenesis and memory formation.

## 2. Results

### 2.1. Specific subpopulation of SUM neurons innervates adult-born neurons

To better understand the connectivity between SUM and abDG neurons, we used a cell-specific trans-synaptic tracing method (*14, 41*) combining retroviral labeling with rabies virus tracing. Specifically, we injected young adult mice with retrovirus expressing the avian TVA receptor, rabies glycoprotein (gp), and green fluorescent protein (GFP) (Rv-Syn-GTRgp) to label dividing DG progenitor cells that develop into abDG neurons over time. After an interval of 4-6 weeks, avian EnvA pseudotyped rabies virus (EnvA-ΔG-MCh) was injected into the DG to trace the afferent inputs of the abDG neurons. (*9, 10, 12–14, 42*) (Figures 1A-D).

**Figure 1.**
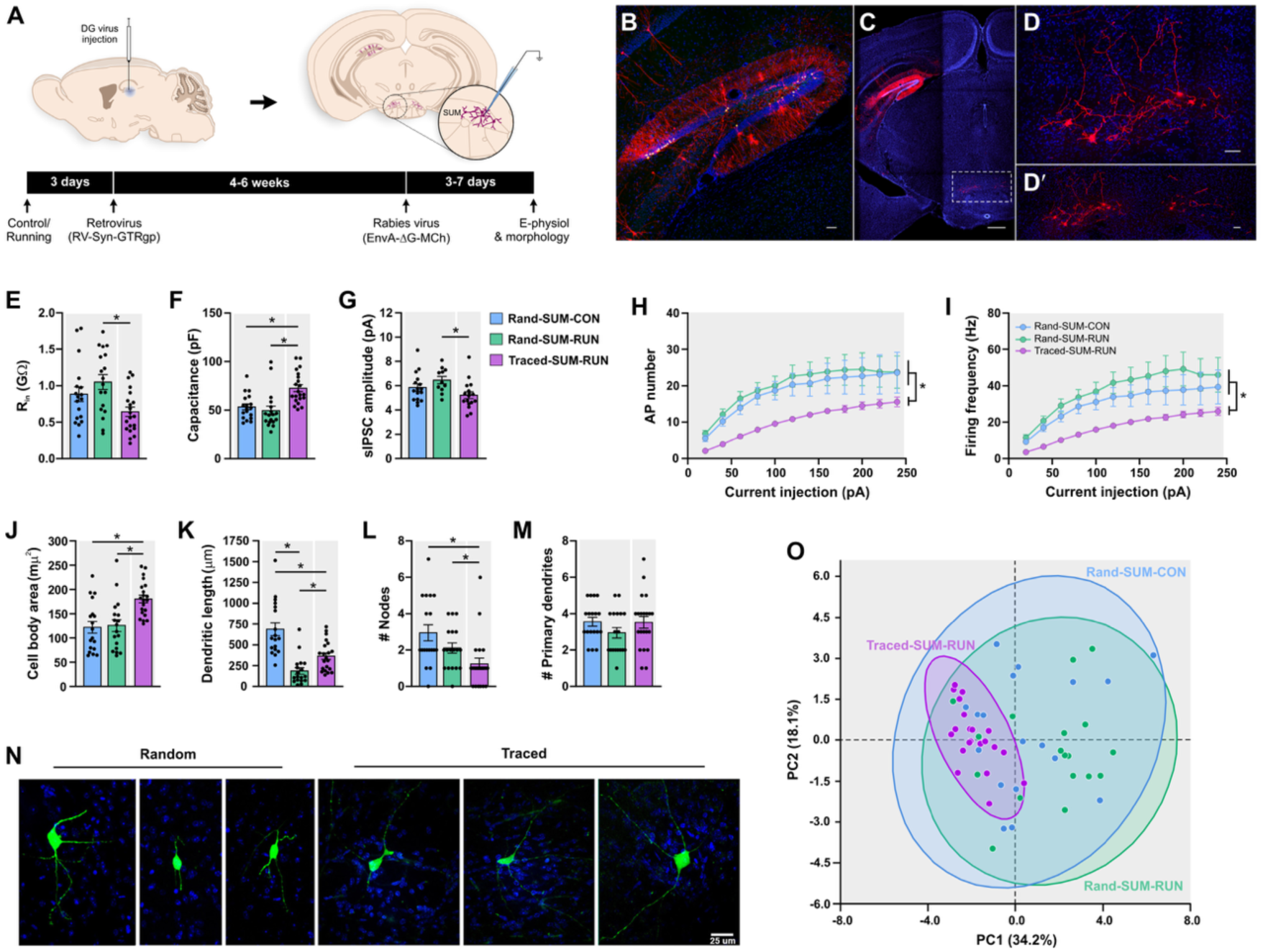
**A subpopulation of SUM neurons innervates adult-born DG neurons. (**A) Timeline of the experiment to label and record supramammillary nucleus (SUM) neurons innervating adult-born neurons in the dentate gyrus (DG). (B-D) Photomicrograph of a coronal section (C) showing the expression of nuclear GFP in retrovirally labeled (RV-Syn-GTRgp) adult-born DG neurons (green) co-expressing mCherry (MCh-GFP, yellow) or MCh (red) only in the SUM (boxed area) following injection of rabies virus (EnvA-ΔG-MCh). High-power photomicrographs of the DG (B) and SUM (D, D’). (E-M) Graphs of the electrophysiological and morphological properties of traced and random SUM neurons from control and runner mice. (E-I) Statistical differences were observed between the mean of traced and random cells in (E) input resistance (R_in_) (F_(2,54)_ = 5.595, *p* = 0.0062) and (F) capacitance (H_(2)_ = 20.32, *p* < 0.0001). (G) The amplitude of the spontaneous inhibitory postsynaptic currents (sIPSC) differs between traced and random groups (H_(2)_ = 8.852, *p* = 0.0120). (H) The number of action potentials (AP) and firing frequency (I) elicited at different current steps were lower in traced than random cells (AP number, F_(2,54)_ = 5.454, *p* = 0.0070; firing frequency, F_(2,54)_ = 6.211, *p* = 0.0037). (J) Traced SUM cells exhibited a larger cell body area than random SUM cells (H_(2)_ = 15.38, *p* = 0.0005). (K) Dendritic length differs between groups (H_(2)_ = 27.27, *p* < 0.0001), with SUM cells having an intermediate length compared to random SUM cells. (L) A smaller number of nodes was observed in traced cells as compared to random cells (H_(2)_ = 13.49, *p* = 0.0012). (M) Random and traced SUM cells have a similar number of primary dendrites (F_(2,54)_ = 1.327, *p* = 0.2738). (N) Biocytin-filled random and traced SUM neurons (green). (O) Principal component analysis (PCA) scatter plot in the 2-dimensional PCA based on all the electrophysiological and morphological properties of the random and traced SUM neurons. PC1 and PC2 captured 52.3% of the total variance. Each dot represents SUM neurons from Rand-SUM-CON (blue), Rand-SUM-RUN (green), and Traced-SUM-RUN (pink) groups. Shaded circles represent 95% confidence regions. Data are means ± SEM.* *p* < 0.05. Statistical analysis was performed using One-way ANOVA (E, M) and Two-way ANOVA RM (H, I), followed by *post hoc* Tukey’s test, as well as Kruskal-Wallis’s test, followed by *post hoc* Dunn’s test (F, G, J, K, L). Rand-SUM-CON, *n* = 18; Rand-SUM-RUN, *n* = 18; Traced-SUM-RUN, *n* = 21. Nuclei stained with DAPI (blue) in B-D and N. Scales bars in B, D, and Dʹ: 50 μm; C: 500 μm and N: 25 μm.

Previously, we have shown that SUM neurons synapse directly onto abDG neurons (*14*) and that the number of trans-synaptically labeled neurons in SUM is substantially increased by running (*13*). Because the SUM afferent input is sparse under sedentary conditions, we performed patch-clamp recordings of traced MCh^+^ SUM neurons in mice housed with a running wheel (Traced-SUM-RUN). Additionally, we recorded random SUM neurons from control (Rand-SUM-CON) and runner (Rand-SUM-RUN) mice. Evaluation of the passive and active membrane properties showed that Traced-SUM-RUN neurons have unique properties, including a lower input resistance (R_in_) and higher capacitance, and exhibit lower action potential (AP) number and firing frequency compared to random SUM neurons from control and runner mice (Figures 1E, 1F, 1H, and 1I). No differences were observed in the resting membrane potential (RMP), time constant (Tau), AP amplitude, and AP half-width (Figures S1A-S1D). Additionally, the Traced-SUM-RUN neurons exhibited less inhibitory control, as the amplitude of spontaneous inhibitory postsynaptic currents (sIPSC) was smaller compared to random SUM neurons (Figure 1G), suggesting that they may be easily excitable. No differences in sIPSC frequency and amplitude and frequency of spontaneous excitatory postsynaptic currents (sEPSC) were observed (Figures S1E-S1G). Morphological analysis showed that Traced-SUM-RUN neurons have a bigger cell body area, intermediate dendritic length, and fewer nodes compared to random SUM cells (Figures 1J, 1K, 1L, and 1N). No differences in the number of primary dendrites were observed (Figure 1M).

A principal component analysis (PCA) of the electrophysiological and morphological properties revealed a clear clustering of the Traced-SUM-RUN neurons projecting to abDG neurons (Figure 1O). The electrophysiological properties that best characterized the first two principal components (PC1 and PC2) were related to the number of AP at 40 pA and R_in_, which accounted for 52.3% of the variable contribution (Figure S2). Altogether, our findings demonstrate that the subpopulation of SUM neurons innervating abDG neurons exhibits unique electrophysiological and morphological properties.

### 2.2 SUM neurons innervating adult-born neurons have a GABAergic and glutamatergic phenotype

Previous studies have shown that SUM cells projecting to the DG co-release glutamate and GABA (*19, 30, 34–40*). To determine the specific phenotype of the subpopulation of SUM neurons projecting to abDG neurons, we evaluated the terminals impinging on abDG neurons. To identify the pre- and post-synaptic compartments, we injected an adeno-associated vector encoding mCherry (AAV-CaMKII-ChR2-mCherry) into the SUM and the retroviral vector CAG-GFP into the DG to label dividing cells (Figure 2A). After a one-month post-surgical interval, to allow the DG progenitor cells to develop into abDG neurons (*3*) and receive SUM innervation (*11, 13, 14*), mice were sacrificed, and tissue sections were analyzed for the presence of vesicular GABA transporter (VGAT) and vesicular glutamate transporter 2 (VGluT2) on SUM neuron terminals (Figure 2B). We found VGAT and VGluT2 labeling at the axon terminals from SUM neurons impinging on abDG neurons. Three-dimensional reconstruction of confocal images from quadruple fluorescent immunolabeled DG shows mCherry-labeled SUM axon terminals (red) co-expressing VGluT2 (blue) and VGAT (white) that synapse onto abDG neurons (green) (Figures 2C and 2C’). This indicates that axon terminals from SUM neurons can accumulate both glutamate (VGluT2^+^) and GABA (VGAT^+^) vesicles in the same terminal. We confirm the expression of VGAT in mCherry-labeled SUM axon terminals by immunoelectron microscopy. The mCherry-labeled VGAT terminals establish asymmetric synapses with dendrites and symmetric synapses with the cell body of abDG neurons (Figure 2D). Taken together, these findings show that SUM axon terminals simultaneously establish spatially segregated asymmetric and symmetric synapses onto abDG neurons, reflecting their capacity to release both glutamate and GABA. These findings support the dual inhibitory and excitatory nature of SUM input to the dentate gyrus (*34, 36, 37, 39, 40*).

**Figure 2.**
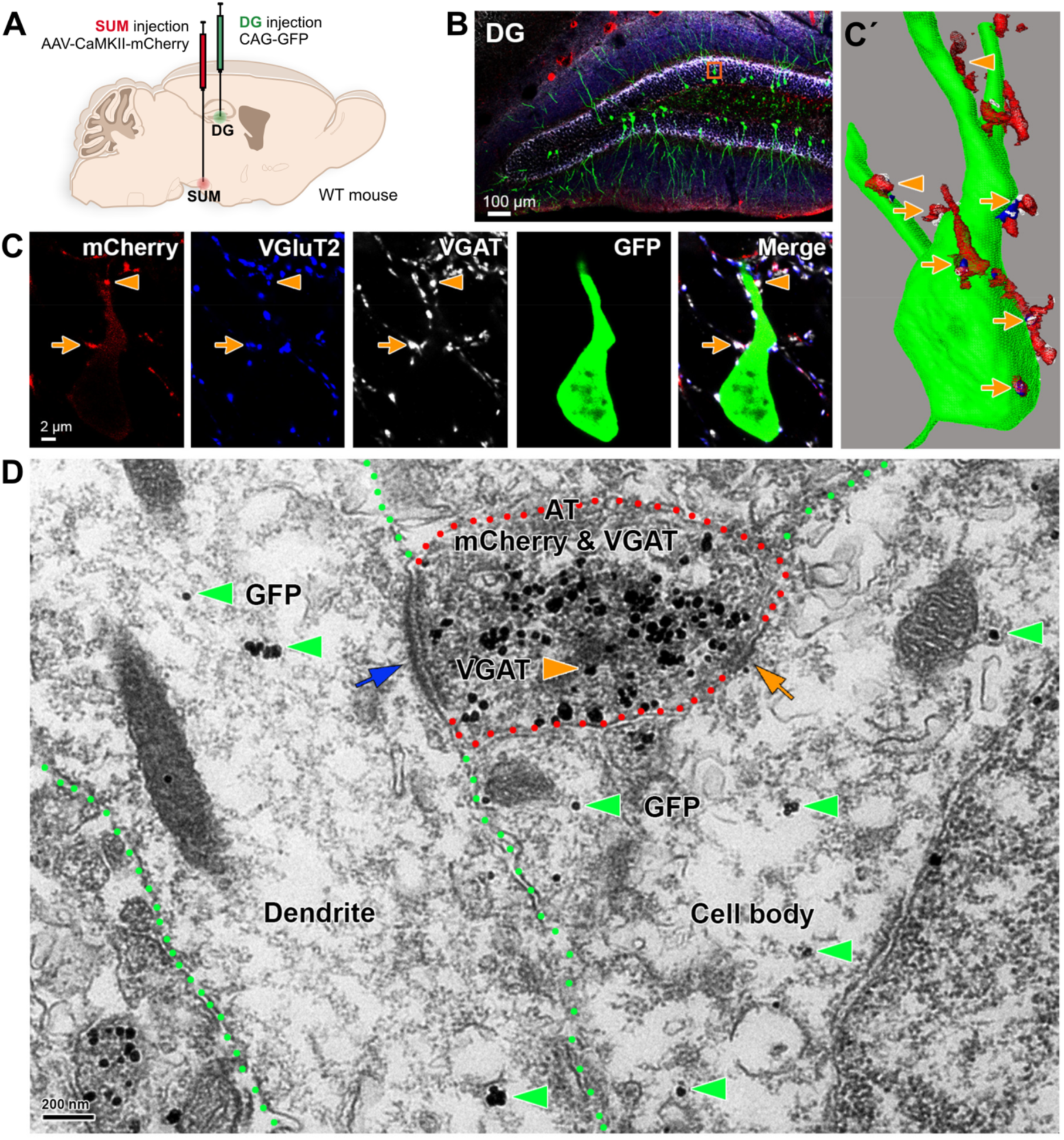
VGluT2^+^/VGAT^+^ SUM terminals make asymmetric and symmetric synapses onto adult-born DG neurons. (A) Experimental design to evaluate the expression of VGluT2 and VGAT in terminals of supramammillary nucleus (SUM) neurons projecting into the dentate gyrus (DG). Viral vector AAV-CaMKII-mCherry was injected into SUM to visualize the axon terminals projecting into the DG. The retrovirus CAG-GFP was injected into the DG to label adult-born DG neurons (abDG). (B) Photomicrograph showing the SUM axon projections expressing mCherry (red), abDG neurons expressing GFP (green), and the immunodetection of VGluT2 (blue) and VGAT (white) in the DG. (C) High magnification of the boxed orange area in (B) showing terminals-like structures co-expressing mCherry, VGluT2, and VGAT, impinging abDG neurons expressing GFP in the cell body (arrows) and dendrites (arrowheads). (C’) Synaptic contacts are better seen after 3D reconstruction. (D) An axon terminal (AT, red outline) from SUM neurons expressing mCherry (dark diffuse material) co-expressing VGAT (orange arrowhead), establishing asymmetric synapses (blue arrow) onto the dendrites (green outline) and symmetric synapses (orange arrow) onto the cell body of abDG neurons expressing GFP (green arrowheads).

### 2.3 Running promotes SUM connectivity and co-release of glutamate and GABA onto adult-born neurons

To determine how the glutamatergic and GABAergic neurotransmission from SUM influences abDG neurons, we injected AAV-hSyn-hChR2-eYFP into SUM and retrovirus CAG-RFP into the DG, followed by whole-cell patch-clamp recordings one month later (Figures 3A and 3B). Specifically, synaptic responses evoked by optogenetic stimulation of SUM terminals were recorded in abDG and developmentally born mature granule cells (mGCs) (*43*) from slices derived from mice housed under control sedentary or running conditions.

**Figure 3.**
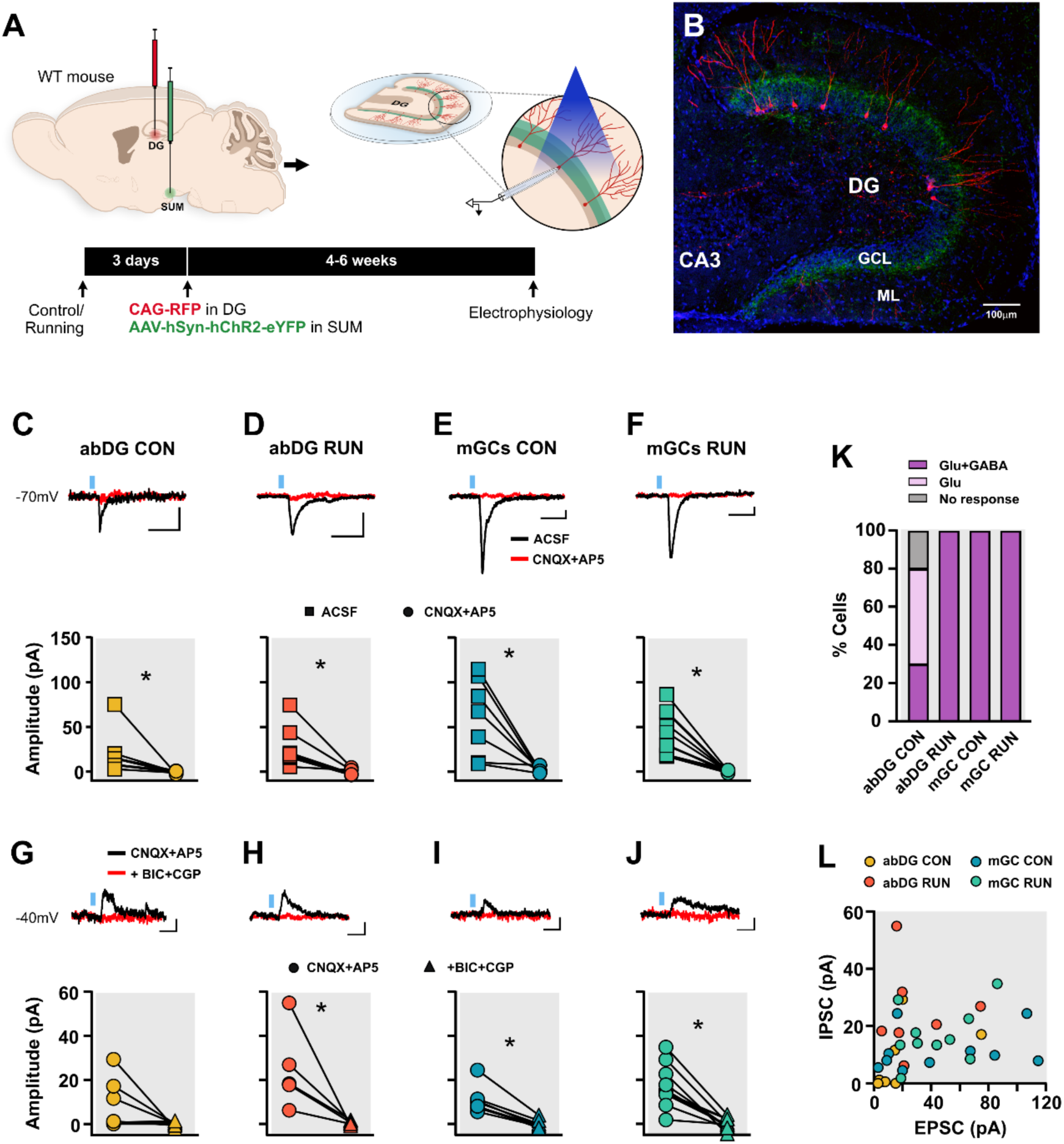
SUM neurons co-release glutamate and GABA onto abDG neurons and mature GCs. (A) Timeline of the experiment. To label dividing cells, control (CON) and runner (RUN) mice were injected with retrovirus expressing red fluorescent protein (CAG-RFP) into the dentate gyrus (DG), whereas to label the SUM projections to the DG, AAV expressing channelrhodopsin and yellow fluorescent protein (AAV-hSyn-hChR2-e-YFP) was injected in the SUM. Four to 6-weeks later, patch-clamp recordings from adult-born granule cells (abDG neurons; RFP^+^) or mature granule cells (mGCs; RFP^-^) were performed in acute hippocampal slices derived from CON and RUN mice. Synaptic responses were evoked by pulses of blue light (465 nm, 1ms). (B) Photomicrograph of a horizontal hippocampal slice showing retrovirally labeled abDG neurons (RFP^+^) and ChR2 expressing SUM axons (YFP^+^, green) in the granule cell layer (GCL) of the DG. Nuclei were labeled with DAPI (blue). (C-F) Upper: Representative traces of inward synaptic currents (Vh = −70 mV) evoked by blue light stimulation of SUM afferents under artificial cerebrospinal fluid (ACSF) conditions (black traces) from (C) abDG CON, (D) abDG RUN, (E) mGCs CON, and (F) mGCs RUN cells. Inward current was blocked by AMPA (10 µm CNQX) and NMDA (100 µm AP5) receptor antagonists in all the groups (red traces). Bottom: The amplitude of the SUM light-evoked inward current (ACSF) was significantly reduced by CNQX+AP5 in (C) abDG CON (W = 36, *p* = 0.0078), (D) abDG RUN (t_(6)_ = 3.552, *p* = 0.012), (E) mGCs CON (t_(6)_ = 3.558, *p* = 0.0115), and (F) mGCs RUN (t_(9)_ = 5.593, *p* = 0.0003) cells. (G-J) Upper: A residual SUM light-evoked outward synaptic current (Vh = −40 mV) was observed after the blocking of glutamatergic receptors (CNQX+AP5; black traces). It was blocked by GABA_A_ [100 µm bicuculine (BIC)] and GABA_B_ [1 µm CGP55845 (CGP)] receptor antagonists in all the cell types (red traces). Bottom: The amplitude of the SUM light-evoked outward current (CNQX+AP5) was significantly reduced by GABA receptors antagonist (BIC+CGP) in (H) abDG RUN (t_(4)_ = 2.859, *p* = 0.046), (I) mGCs CON (W = −28, *p* = 0.0156), and (J) mGCs RUN (t_(9)_ = 5.834, *p* = 0.0002), but not in all the (G) abDG CON (W = −13, *p* = 0.1250). (K) Percentages of cells exhibiting both glutamatergic and GABAergic (Glu+GABA), only glutamatergic (Glu), or no response by blue light stimulation of SUM afferents. (L) Scatter plot showing the relationship between inward EPSC and outward IPSC amplitude in abDG neurons and mGCs from CON and RUN mice. No significant correlation was observed across any of the groups. abDG CON (r = 0.5007, *p* = 0.9250), abDG RUN (r = −0.01966, *p* = 0.9666), mGCs CON (r = 0.2624, *p* = 0.4640), mGCs RUN (r = 0.3980, *p* = 0.2547). Data are means ± SEM.* *p* < 0.05. Statistical analysis was performed using paired t-test (D, E, F, H, J), Wilcoxon paired test (C, I, G), and Pearson correlation coefficient (K).

First, we confirmed channelrhodopsin (ChR2) expression in SUM neurons by whole-cell patch-clamp recordings from YFP^+^ cells located in brain slices containing SUM. We delivered 465 nm light of 1 ms pulse width at 1 Hz, which reliably evoked action potentials in ChR2-eYFP^+^ SUM neurons (Figure S3). Next, SUM axons innervating the granule cell/inner molecular layers of DG were optically stimulated in hippocampal brain slices while recording from either abDG neurons or mGCs. To examine the co-release of glutamate and GABA from the SUM terminals, we recorded optically evoked postsynaptic currents. In the presence of artificial cerebrospinal fluid (ACSF) (*Vh* = −70 mV), blue light pulses evoked an inward current that was completely blocked by AMPA and NMDA receptor antagonists, 6-Cyano-7-nitroquinoxaline-2,3-dione (CNQX) and DL-2-amino-5-phosphonovalerate (DL-AP5), respectively, in both abDG neurons and mGCs from control and runner mice (Figures 3C-F). This indicates that the SUM terminals release glutamate, eliciting EPSCs. A residual outward current was observed at *Vh* −40 mV in both abDG neurons and mGCs from both groups, which was completely blocked by GABA_A_ and GABA_B_ receptor antagonists bicuculline (BIC) and CGP55845 (CGP), respectively (Figures 3G-3J). This indicates that SUM terminals also release GABA, eliciting IPSCs. Quantitatively, we found that in 10 abDG neurons recorded from control mice, light stimulation elicited both EPSCs and IPSCs in 3 cells (30%), EPSCs only in 5 cells (50%), and no response in 2 cells (20%) (Figure 3K). In contrast, voluntary running promoted GABA release from SUM terminals, with light stimulation eliciting both EPSCs and IPSCs in 100% of the recorded abDG neurons—a response pattern similar to that observed in mGCs from control and runner mice (Figure 3K). These findings are in agreement with the increased connectivity between SUM and abDG neurons induced by runner mice (*13, 16*) and studies suggesting exercise promotes GABA release (*44, 45*). Next, we examined the relationship between excitatory and inhibitory inputs by analyzing the correlation between EPSC and IPSC amplitudes in individual abDG neurons and mGCs. There was no significant correlation in either cell type across both CON and RUN mice (Figure 3L), suggesting that glutamate and GABA are released from independent vesicles (*46*). Altogether, these findings show that voluntary running strengthens SUM synaptic connectivity and boosts GABA release onto abDG neurons, suggesting a potential functional role for the SUM-GABAergic pathway in cognitive behavior.

### 2.4 Chemogenetic silencing of SUM GABAergic neurons impairs pattern separation behavior

To determine the functional relevance of the SUM-GABA–DG circuit, we investigated whether the selective silencing of SUM-GABAergic inputs alters pattern separation performance. Male VGAT-ires-Cre mice were injected into the SUM with pAAV2-hSyn-DIO-hM4D(Gi)-mCherry to silence SUM-to-DG GABAergic transmission (Figures 4A and 4B). Behavioral training and testing commenced four weeks post-surgery. Following touchscreen pre-training shaping protocols, mice underwent task training in which they discriminated between visually identical, spatially discrete stimuli separated by an intermediate distance to obtain a liquid reward. After completion of task training, mice underwent two rounds of 16-day probe trials presenting either small or big separations between the stimuli. Mice were injected with either saline or Clozapine N-oxide (CNO) daily, 30 min before the onset of testing. For each session, mice had to reach a criterion of seven out of eight correct trials before the stimulus positions were reversed. Mice that received saline during the first 16 days were injected with CNO during the second 16 days, and vice versa (Figure 4A). Analysis of the number of trials to criterion revealed significant main effects of both stimulus separation and treatment. Specific comparisons showed that CNO-treated mice required significantly more trials to reach criterion than saline-treated mice during small separation, whereas no significant difference was observed during big separation (Figure 4C). The analysis of the total number of reversals completed across sessions showed that CNO mice had significantly fewer reversals than saline controls during both the small and big separation sessions (Figure 4D).

**Figure 4.**
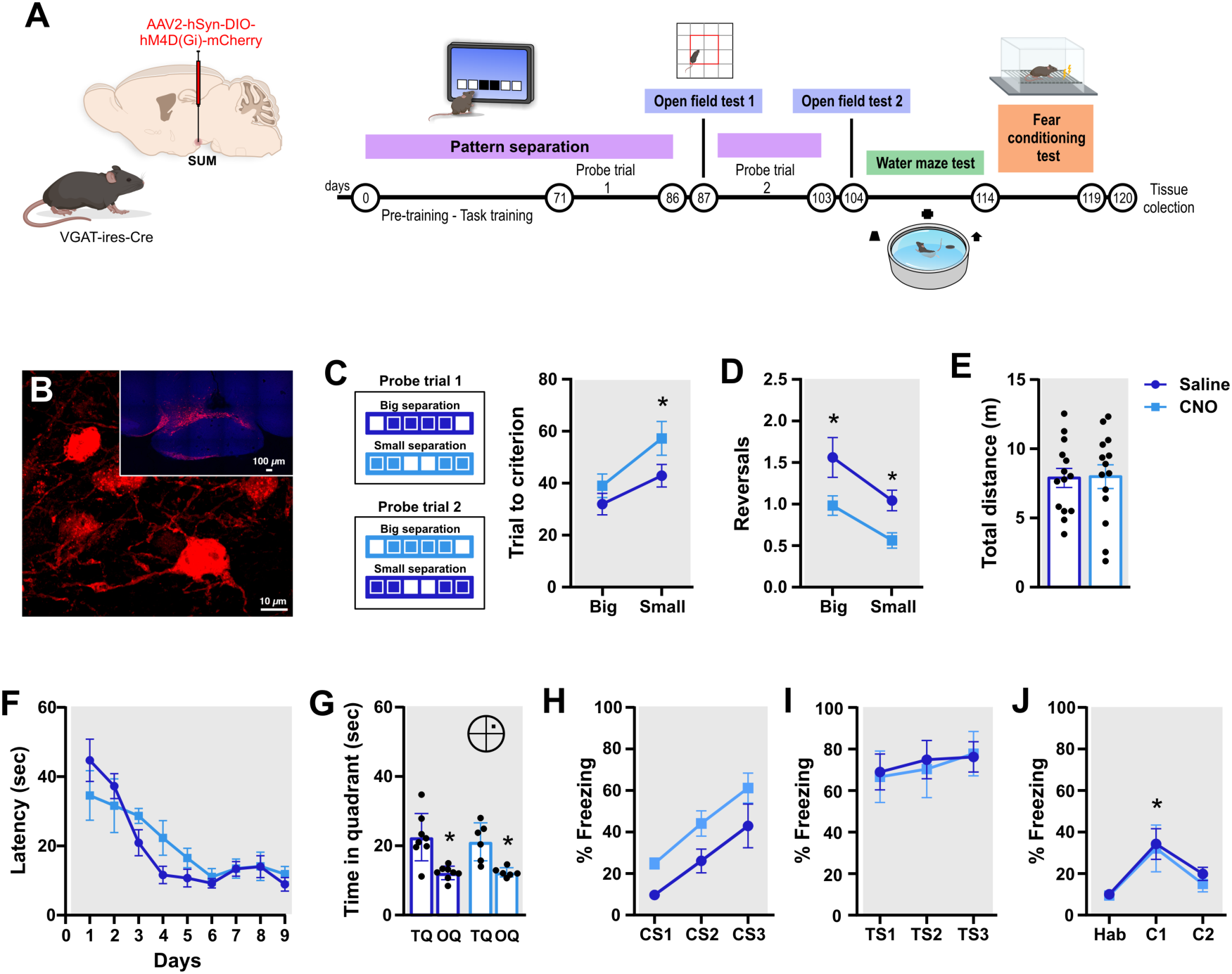
Pattern separation is impaired by inhibition of SUM-projecting GABAergic neurons. (A) Timeline of the experiment. Mice underwent stereotaxic surgery to inject pAAV2-hSyn-DIO-hM4D(Gi)-mCherry into the supramammillary (SUM) nucleus. Four weeks thereafter, mice started habituation and pre-training in the touchscreen task (duration 71 days). Next, mice were divided into control saline and CNO groups for two rounds of 16-day probe trials (Probe trial 1). Mice who received CNO in probe 1 subsequently received saline, and mice who received SAL subsequently received CNO for the second set of probe trials (Probe trial 2). Following completion of each set of probe trials, mice were evaluated in the open field test following saline or CNO injection. Thereafter, mice were trained and tested in the Morris water maze and fear conditioning paradigms following daily saline or CNO injection before training and testing. Upon completion of behavioral testing, mice were deeply anesthetized and perfused for histological analysis. (B) Photomicrograph of MCh^+^ cells in the SUM, overview and high magnification image in a tissue section derived from an AAV-injected VGAT-ires-cre mouse. Nuclei are stained with DAPI. (C) The touchscreen paradigm acquisition of big and small separation between two identical stimuli was tested in saline- and CNO-treated mice, which were trained to reach a criterion where seven of eight trials were performed correctly. Pattern separation differed between the groups as evidenced by main effects of separation (F_(1,13)_ = 17.80, *p* = 0.001) and treatment (F_(1,13)_ = 10.01, *p* = 0.0075), albeit no interaction (F_(1,13)_ = 1.540, *p* = 0.23). Compared to saline controls, CNO-treated mice exhibited significantly impaired performance when the separation between the stimuli was small (*p* = 0.019), but not when the separation between stimuli was big (*p* = 0.36). (D) Reversal of stimulus reinforcement in the big and small conditions. The reinforcement of the stimuli was reversed each time a mouse reached criterion so that the previously correct location became the incorrect location and vice versa. CNO administration significantly reduced the number of reversals for big and small separations, as evidenced by a main effect of separation (F_(1,13)_ = 19.88, *p* = 0.0006) and treatment (F_(1,13)_ = 16.32, *p* = 0.0014), albeit no interaction (F_(1,13)_ = 0.39, *p* = 0.53). (E) There was no difference between the groups in total distance travelled in the open field (t_(13)_ = 0.23, *p* = 0.82). (F) Morris water maze learning over 9 days with 4 trials per day did not show a difference between groups (treatment effect: F_(1,12)_ = 0.3921, *p* = 0.5429). (G) Retention of spatial memory in the probe trial conducted 24 h after the last training session did not differ between groups (treatment effect: F_(1,12)_ = 0.12, *p* = 0.73). TQ, target quadrant; OQ, other quadrants. (H) In the fear conditioning paradigm, freezing was similar between groups in the conditioning session (treatment x sessions: F_(2,24)_ = 0.710, *p* = 0.9317) and increased over tone-shock (CS1, CS2, CS3) pairings (F_(2,24)_ = 29.30, p < 0.0001). (I) the tone-cued (TS1,TS2,TS3) phase (treatment x session: F_(2,24)_ = 0.1515, *p* = 0.8603). (J) Both groups displayed increased freezing to context (C), (session effect: F_(2,24)_ = 13.79, *p* = 0.0001), as compared to habituation (Hab) on day 3 (*p* < 0.05). Data are means ± SEM. * *p* < 0.05. Statistical analysis was performed using Two-way ANOVA RM (C, D, F, G, H, I and J), followed by *post hoc* Tukey’s test, and paired t-test (E).

#### Open Field

To determine whether CNO treatment altered general locomotor activity or anxiety-like behavior, which could confound learning performance, mice were evaluated in the open field after each round of probe trials under saline or CNO conditions. The total distance traveled did not differ between saline and CNO conditions (Figure 4E). Furthermore, no significant differences were observed between groups regarding the distance traveled or time in either the center or periphery of the open field (Figures S4A and S4B), showing that CNO administration did not alter locomotor activity or anxiety-like behavior.

#### Morris water maze and fear conditioning tests

To examine whether the deficit in pattern separation was due to a more general memory dysfunction, we tested the mice in the Morris water maze and contextual fear conditioning paradigm. We did not observe a difference between the saline and CNO groups in acquisition of the water maze training over nine days with four trials per day, as evidenced by similar latencies to the platform (Figure 4F). In the probe trial, 24 hr after the last training session, both groups spent more time in the platform quadrant compared to the other quadrants, indicating retention of spatial memory (Figure 4G). Swim speed (Saline, 15.08 ± 0.48 cm/sec; CNO, 15.22 ± 0.80 cm/sec, *p* = 0.87) also did not differ between the groups.

Fear learning and memory were assessed by measuring the degree of freezing elicited by pairing a tone with a shock. In the habituation (Hab) session on Day 1, there were no differences in freezing between the groups (Saline, 10.02 ± 2.18%; CNO, 9.49 ± 1.84%; *p* = 0.85). On Day 2, the Conditioning Phase consisted of three tone-shock pairings. Both groups displayed an increase in freezing during conditioning (Figure 4H). On Day 3, mice were tested for tone-cued memory session in a new contextual environment (Figure 4I). One hour after the tone-cued session on day 3, and on day 5, mice were tested in the original environment used in the habituation (Hab) and conditioning sessions to evaluate contextual memory. There was no difference between the saline and CNO groups (Figure 4J).

Furthermore, to evaluate whether inhibiting GABAergic SUM input to the DG affects granule cell activation, mice were exposed to a novel cage with a running wheel for one hour before euthanasia. The number of c-Fos^+^ cells in the granule cell layer was reduced (Figures S4C-S4E).

## Discussion

In the present study, we characterized the physiological properties and functional contribution of SUM neurons projecting to adult-born neurons. Patch-clamp recordings revealed a SUM subpopulation with unique electrophysiological properties. These neurons exhibit lower intrinsic excitability and a larger soma, whereas the general SUM neuronal population includes high-spiking cells that do not impinge on newborn neurons. Electron microscopy and patch-clamp recordings demonstrated that SUM neurons form both symmetric (putative inhibitory) and asymmetric (putative excitatory) synapses onto abDG neurons, which co-release GABA and glutamate. Furthermore, voluntary wheel running enhanced SUM GABAergic transmission onto abDG neurons, whereas chemogenetic silencing of GABAergic release impaired pattern separation but did not affect spatial memory or fear conditioning. Together, our findings indicate that a discrete subset of SUM neurons provides input to new neurons and modulates pattern separation behavior via inhibitory neurotransmission.

To selectively identify SUM neurons that provide input to abDG neurons, we utilized retroviral labeling of abDG neurons combined with rabies virus-mediated retrograde monosynaptic trans-synaptic tracing (*12–14, 47*). While sparse SUM projections to abDG neurons preclude robust electrophysiological characterization under sedentary control conditions, voluntary running substantially strengthens this connectivity, enabling their characterization. By recording traced SUM neurons from runner mice *in vitro*, we identified a unique population of SUM neurons with distinctive morphological and electrophysiological properties that innervate new neurons. Specifically, SUM neurons projecting to abDG neurons have a larger soma, which correlates with their biophysical properties, including reduced input resistance, higher membrane capacitance, and lower intrinsic excitability, compared with the general SUM neuron population, regardless of whether the recordings from this general population were made from cells derived from mice housed under sedentary or running conditions. Indeed, the electrophysiological properties of SUM cells that innervate new neurons differ from the subpopulations of SUM neurons projecting to the dorsal and ventral DG (*48*) and SUM^tac1+^ projecting neurons associated with locomotion (*49*). In contrast to those previous studies that focused on a limited number of properties (e.g., firing rate, AP frequency), our findings reveal that the identified cells diverge significantly across multiple physiological and morphological parameters. Specifically, measurements of capacitance, input resistance, firing rate, and soma size indicate that SUM neurons projecting to abDG neurons are larger and have a unique electrophysiological profile that has not been previously characterized.

Subpopulations of SUM neurons targeting the hippocampus express vesicular transporter markers of both GABA and glutamate release at the axon terminals (*37, 38, 50*). Indeed, optogenetic stimulation of SUM terminals expressing VGAT and VGluT2 elicits monosynaptic release of both glutamate and GABA onto dentate granule cells (*36, 38, 39*). Consistent with these studies, we show the co-expression of VGluT2 and VGAT in terminals of the unique population of SUM neurons that project to new neurons. Such axonal terminals were preferentially located around the cell body and proximal dendrites of abDG neurons, forming segregated symmetrical (putative inhibitory) and asymmetrical (putative excitatory) synapses, reflecting their capacity to release both glutamate and GABA. Optogenetic activation of SUM neurons confirmed glutamatergic and GABAergic co-transmission onto new and mature DG neurons. However, only a third of adult-born neurons from sedentary control mice exhibited this response; the remaining cells were either glutamatergic only or did not respond. A previous report describes GABAergic responses that precede glutamatergic neurotransmission in very young two-week-old abDG neurons (*36*). However, our tracing studies have shown that, like entorhinal cortex input, SUM innervation arrives later in the development of abDG neurons, during the critical period of heightened plasticity (*16, 42*). Interestingly, voluntary running promoted GABA release from SUM terminals onto one-month-old neurons, similar to previously observed effects of exercise on mature DG granule neurons (*19, 37–39*), cortex and other brain areas (*44, 51, 52*). Indeed, all adult-born neurons derived from runner mice responded with both glutamatergic and GABAergic receptor currents upon SUM terminal optogenetic stimulation. Enhancement of GABAergic transmission onto the DG potentially controls spike timing, facilitating the temporal coupling required for memory discrimination (*53*). The increased GABAergic inhibition onto adult-born neurons may also enhance pattern separation by promoting sparse and transient neuronal spiking, supporting the role of adult neurogenesis in preventing overlap between similar memories (*6, 54–57*).

To obtain further insight into the functional role of GABAergic neurotransmission in this pathway, we tested whether chemogenetic silencing of SUM GABA affects hippocampus-dependent behaviors. Specifically, mice were tested for pattern separation ability in the touchscreen, spatial memory in the Morris water maze and contextual fear conditioning. Only pattern separation was affected by chemogenetic inhibition of SUM GABAergic neurotransmission. During probe trials, the number of trials to reach criterion was increased when the distance between identical stimuli was small. In addition, we observed fewer reversals of stimulus location in both small and large separation conditions. The ability to make fine discriminations has been associated with the functional role of adult-born neurons in pattern separation (*6, 14, 22, 23, 54*) suggesting that SUM neurons innervating adult-born neurons play an important role in episodic-like memory. Overall, SUM inhibitory neurotransmission may affect sparse encoding by DG granule cells and thereby pattern separation ability.

We did not observe changes in spatial memory or fear conditioning, suggesting that SUM GABAergic transmission plays a selective role in pattern separation. In the Morris water maze, there was no difference between the groups in acquisition or retention of spatial memory. In previous studies, lidocaine-reversible inactivation of the SUM, before the training and probe trials in the Morris water maze, also did not affect performance (*33, 58*). However, when SUM was transiently inactivated after training trials, retention of spatial memory was impaired (*33*). It has also been reported that SUM silencing reduces contextual fear memory (*48, 58*). However, we observed no changes in contextual or tone-cued fear memory. Together, these findings indicate that SUM modulation of spatial memory and fear conditioning may rely on glutamatergic neurotransmission, whereas pattern separation behavior is specifically mediated by GABAergic signaling.

In this study, we have identified a discrete population of SUM neurons that exhibit unique morphological and electrophysiological properties and co-release glutamate and GABA onto abDG neurons. Furthermore, we demonstrate that chemogenetic silencing of GABA transmission from SUM neurons, which includes this selective population projecting SUM-abDG neurons, impairs pattern separation. Together, these findings identify a novel pathway in synaptic plasticity and memory function.

## Experimental Procedures

### Subjects

Male C57Bl/6J mice (Jackson Labs #000664) designated for electrophysiological experiments were individually housed at two months of age and randomly assigned to control or voluntary wheel-running conditions. Mice were housed in a 12h light–dark cycle (lights on at 6:00 a.m. and off at 6:00 p.m.) with food and water *ad libitum*. The experimental protocols were approved by the National Institute on Aging Institutional Animal Care and Use Committee (IACUC).

Adult male VGAT-ires-Cre mice (Jackson Labs #028862) were utilized for behavioral experiments. Mice were 13-15 weeks old (n = 35) and group-housed (3-5 per cage) in standard cages with a 12h light–dark cycle (lights on at 7:30 a.m. and off at 7:30 p.m.) with water and food *ad libitum*. Florida Atlantic University’s IACUC approved all experimental procedures. All procedures were conducted in accordance with the guidelines required by the National Institutes of Health Guide for the Care and Use of Laboratory Animals.

### Viral vectors

Retroviral vector RV-SYN-GTRgp, expressing nuclear green fluorescent protein (GFP), avian TVA receptor, and rabies virus glycoprotein (Rgp) driven by the neuron-specific synapsin promoter, retroviral vector CAG-GFP, retroviral vector CAG-RFP, and EnvA-pseudotyped rabies virus (EnvA-ΔG-MCh) were produced as previously described (*14, 41, 59*) and injected intracerebrally as detailed in the Supporting Text.

### Brain slice preparation and in vitro electrophysiological recordings

Animals were deeply anesthetized with isoflurane and then rapidly decapitated. Brains were transferred to cooled (1–4 °C) dissection solution, containing (mM): 110 Choline-Cl, 2.5 KCl, 1.25 NaH_2_PO_4_, 25 NaHCO_3_, 25 glucose, 1 CaCl_2_, 7 MgCl_2_, 0.6 Na^+^ pyruvate, 1.3 Na^+^ ascorbate, and 3 kynurenic acid. Horizontal hippocampal slices (300 μm thick) were obtained and transferred to a chamber containing incubation solution (in mM): 125 NaCl, 2.5 KCl, 1.25 NaH_2_PO_4_, 25 NaHCO_3_, 2 CaCl_2_, 2 MgCl_2_, 20 glucose, 3 Na^+^ pyruvate, and 1.3 Na^+^ ascorbate in distilled water (equilibrated with 95% O_2_ and 5% CO_2_; pH 7.4). Slices were incubated at 34 °C for 10 min and stored at room temperature for >1 hour before patch-clamp experiments were conducted as described in the Supporting Text.

### Electrophysiology data analysis

All current clamp recordings were taken at the resting membrane potential (RMP) of the neuron. RMP was determined during the silent period of neuronal spontaneous activity. A current step protocol ranging from 20 pA to 240 pA of 600 ms duration was applied to determine the action potential firing profile and current injection relationship. Input resistance was determined from the slope of the linear regression fit through the V–I plot constructed by plotting the amplitude of the steady-state voltages (V) against the corresponding step current injections (I) from −10 to +5 pA of 1000 ms duration with 5 pA steps. Data analysis is described in the Supporting Text.

### Optically evoked synaptic stimulation

For optical stimulation, light was produced by a 465-nm Plexibright LD-1 LED module (Plexon, Dallas, TX, USA) and delivered by an optical fiber (200 μm core diameter, NA = 0.5) that was submerged in ACSF and placed ∼300 μm from the recording site. SUM axons expressing ChR2 were activated with the optical fiber placed just above the DG granule cell layer as described in the Supporting Text.

### Electron microscopy and immunohistochemistry

Methods were described previously (*60*). Briefly, vibratome tissue sections from three male WT-ChR2-mCherry-GFP mice (C57BL/6 mice injected with AAV5-CaMKII-ChR2-mCherry in SUM and CAG-GFP in DG, respectively; Fig. 2A) were rinsed and incubated with 1% sodium borohydride to inactivate free aldehyde groups, rinsed, and then incubated with blocking solution. Tissue processing and ultrastructural analysis of brain tissue is further described in the Supporting Text.

### Behavioral assays, General Procedure and Overview

Adult male VGAT-ires-cre mice (Jackson Labs #028862), 13-15 weeks old at the beginning of the experiment (n = 35) were group-housed (3-5 per cage) in standard cages with *ad libitum* access to food and water on a standard 12h/12h day/night cycle: lights were switched on at 7:30 AM and off at 7:30 PM. Mice (n = 35) underwent stereotaxic surgery to inject pAAV2-hSyn-DIO-hM4D(Gi)-mCherry into the SUM nucleus, resulting in the attrition of 5 mice. Four weeks thereafter, mice (n = 30) started habituation and pre-training in the touchscreen task. Mice were food-restricted from 8 days to ∼4 grams of chow per day, starting before the beginning of the touchscreen experiment. Exclusion criteria resulted in n = 22 mice for task training, which was successfully completed by n = 20 mice. These mice were divided into saline control and CNO groups (n = 10 per condition), for two pattern separation probe trial sessions of 16 days each. Mice who received CNO in the first probe trial subsequently received saline, and mice who received saline subsequently received CNO for the second set of probe trials. Following completion of each set of probe trials, mice were evaluated in the open field test following saline or CNO injection. Next, mice were trained in the Morris water maze for 9 days and then tested for memory retention in a probe trial 24 h after the last training trial following saline or CNO injection. Finally, mice were injected with saline or CNO and tested in the contextual fear conditioning paradigm (Figure 4A). Following completion of behavioral testing and before brain tissue collection, mice were injected with saline or CNO and housed in a new cage with a running wheel for one hour for c-Fos activation. Immediately thereafter, mice were deeply anesthetized and perfused transcardially. Histological analysis of the injection site location and spread of the vector resulted in the inclusion of n = 14 mice in the final data analysis.

### Surgery

Mice 13-15 weeks old, underwent stereotaxic surgery. Mice were anesthetized with isoflurane (Abbott) and stereotaxic surgery was performed to deliver 0.2 μl of pAAV2-hSyn-DIO-hM4D(Gi)-mCherry (≥ 5×10¹² vg/mL, Addgene, #44362-AAV2) into the supramammillary nucleus using spatial coordinates relative to Bregma as follows: AP = −2.70 mm; ML = 0.00 mm; DV = −4.90 mm.

### Drug Administration

Clozapine N-oxide (CNO) (freebase) (HelloBio, HB1807) was dissolved in 0.9% saline at 1mg/ml and administered by intraperitoneal (i.p.) injection at 5 mg/kg. CNO or an equivalent volume of 0.9 % saline was administered daily, 30 min prior to the start of probe trial testing for the pattern separation task in the touchscreen, and similarly prior to open field behavior, water maze training and testing, and in the fear conditioning paradigm before habituation, conditioning, tone-cued and context testing.

### Touchscreen-based pattern separation test

The testing apparatus for the location discrimination (LD) task (Lafayette, Cambridge) includes a sound attenuating cubicle (width 56.5 cm × depth 53.2 cm × height 54.5 cm) and a touch screen chamber (width at feeder 4.6 cm × width at screen 23.8 cm × depth 17 cm × height 23 cm) consisting of a black Plexiglas trapezoidal wall, touch screen installed with infrared sensor (WhiskerServer Controller), a perforated-metal floor, reward area, a liquid pump feeder, a speaker with tone generator, a tray light and a house light. A black Plexiglas “mask” panel, which includes two rows of six square windows (2.5 cm × 2.5 cm, equally spaced 1 cm apart), located 1.6 cm from the bottom of mask panel was inserted in front of the touch screen to allow for presentation of white stimuli on the 6 windows at the bottom of the mask panel. When an animal touches the white stimulus coming from one of the 6 windows, the software (ABET II Touch software) counted and saved its activity automatically. The touchscreen training and testing is described in Supporting Text.

### Open field test

Spontaneous locomotor activity was evaluated after the first and second rounds of 16-day probe trials by using an infrared photobeam-controlled open-field testing chamber (Version 4.0, Med Associates, St. Albans, VT, USA; height 20.3 cm, width 27.3 cm, depth 27.3 cm). The center zone was defined as an area of 14.29 cm × 14.29 cm. Mice were injected with either saline or CNO (5 mg/kg) 30 min before they were placed in the open field chamber and explored for 60 min. Mouse movement was detected by three 16-beam infrared arrays installed on X, Y, and Z axes and the time and distance in the center and periphery of the open field were automatically recorded by Activity Monitor software.

### Morris water maze

Spatial learning and retrieval were evaluated using the Morris water maze paradigm (*61*) as described in the Supporting Text.

### Fear Conditioning

Pavlovian conditioning-based approach was used to evaluate fear memory. Mice were tested within a four-chamber system utilizing a near-infrared video conditioning system (MED Associates Inc., Fairfax, VT, USA) that allows for testing of four animals simultaneously. Each chamber consists of stainless-steel walls with one transparent plexiglass door and a floor of parallel stainless-steel rods connected to a shock generator. Fear conditioning is described in the Supporting Text.

### Immediate early gene expression

To validate the use of CNO we quantified c-Fos in the DG granule cell layer. After completion of behavioral testing, animals were given an injection of either saline or CNO (5 mg/kg). Thirty min after the injection, mice were individually housed in a new cage with access to a running wheel for one hour. Immediately thereafter mice were deeply anesthetized and perfused transcardially and tissue was processed for c-Fos analysis as described in the Supporting Text.

### Statistical analysis

Statistical analyses were performed using GraphPad Prism 7 (GraphPad Software Inc, CA). ANOVA was used for multiple comparisons and the Tukey’s test or Fisher’s PLSD test for the post-hoc analysis. Student’s t-test was conducted for normally distributed data. Two-sample Kolmogorov–Smirnov test was conducted to compare population distributions. Averaged data are shown as mean ± SEM and statistical significance was concluded at p < 0.05 for all analysis.

## Acknowledgements

We thank the FAU NeuroBehavioral and Comparative Medicine Cores, and Dr. Jana Boerner of the NeuroImaging Core, Stephanie Vitek for graphic art contributions and Linda R. Kitabayashi photomicrograph composition. Funding: Secihti CF-2023-I-1706 and in part by the Intramural Research Program of the NIH and by the FAU Stiles-Nicholson Brain Institute and the Jupiter Life Sciences Initiative.

