## Supplemental Figures and Methods for "Select properties of supramammillary input to adult-born dentate gyrus neurons"

#### Figure S1

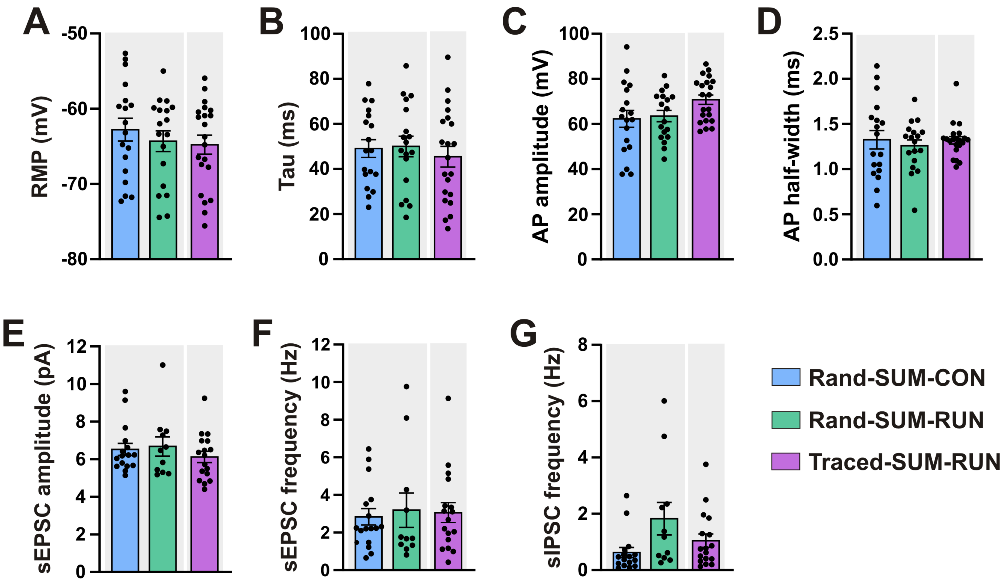


**Fig. S1. Intrinsic and synaptic properties of traced and random SUM cells.** (A-B) Graphs of the passive properties of traced (Traced-SUM-RUN) and random (Rand-SUM-CON and Rand-SUM-RUN) SUM neurons. Traced and random SUM cells exhibit similar (A) resting membrane potential (RMP; $H_{(2)} = 1.237$, $p = 0.5387$) and (B) time constant (Tau; $F_{(2,54)} = 0.3173$, $p = 0.7294$). (C-D) Graphs of the action potential (AP) properties of traced and random SUM neurons. No significant differences were observed in the (C) AP amplitude ($F_{(2,54)} = 2.757$, $p = 0.0724$) and (D) AP half-width ($F_{(2,54)} = 0.2543$, $p = 0.7764$) between traced and random SUM neurons. (E-F) Graphs of the mean (E) amplitude and (F) frequency of the spontaneous excitatory postsynaptic currents (sEPSC). Traced and random SUM neurons receive similar glutamatergic inputs (sEPSC amplitude, $H_{(2)} = 0.9173$, $p = 0.6321$; sEPSC frequency, $H_{(2)} = 0.3208$, $p = 0.8518$). (G) The mean frequency of the spontaneous inhibitory postsynaptic current (sIPSC) is similar in traced and random SUM neurons ($H_{(2)} = 5.561$, $p = 0.062$). Data are means ± SEM. Statistical analysis was performed using One-way ANOVA (B, C, D) followed by *post hoc* Tukey's test, and Kruskal-Wallis's test followed by *post hoc* Dunn's test (A, E, F, G).

### Figure S2

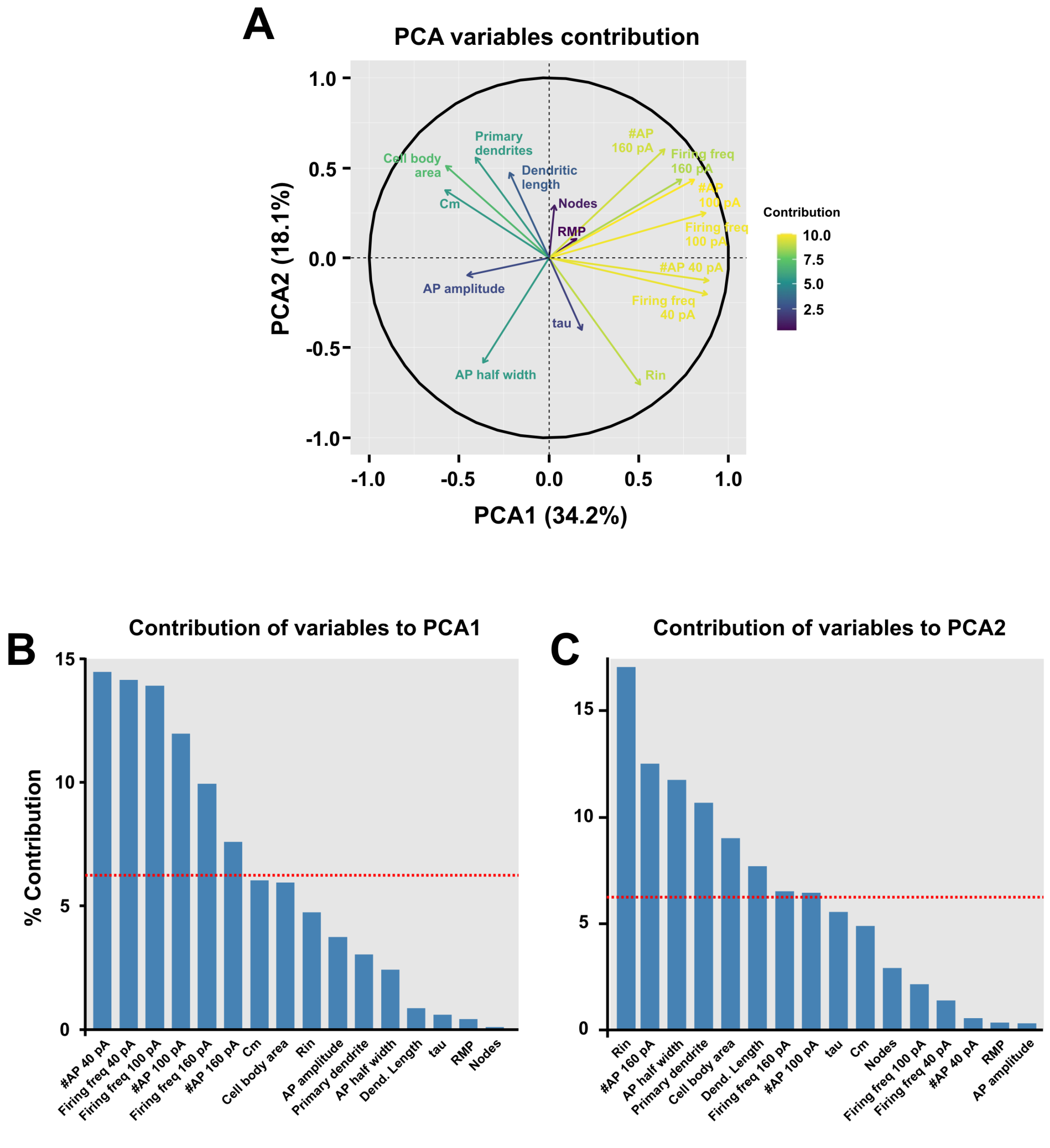


**Fig. S2. Principal component analysis (PCA).** (A) PCA of variables correlation plot in the 2-dimensional PCA based on all the electrophysiological and morphological properties of the random and traced SUM neurons. PC1 and PC2 captured 52.3% of the total variance. The arrows represent the variables. When arrows are far from the center and close to each other, they are positively correlated, whereas when they are symmetrically opposed, they are negatively correlated. If the arrows are orthogonal, they are not correlated. Color represents the contribution of the variables. B-C) Bar plot displaying the percentage of contribution of each of the electrophysiological and morphological properties of the traced (Traced-SUM-RUN) and random (Rand-SUM-CON and Rand-SUM-RUN) SUM neurons to (B) PCA1 and (C) PCA2. The red dashed line on the graphs (B and C) indicates the average contribution (7.69%).

### Figure S3

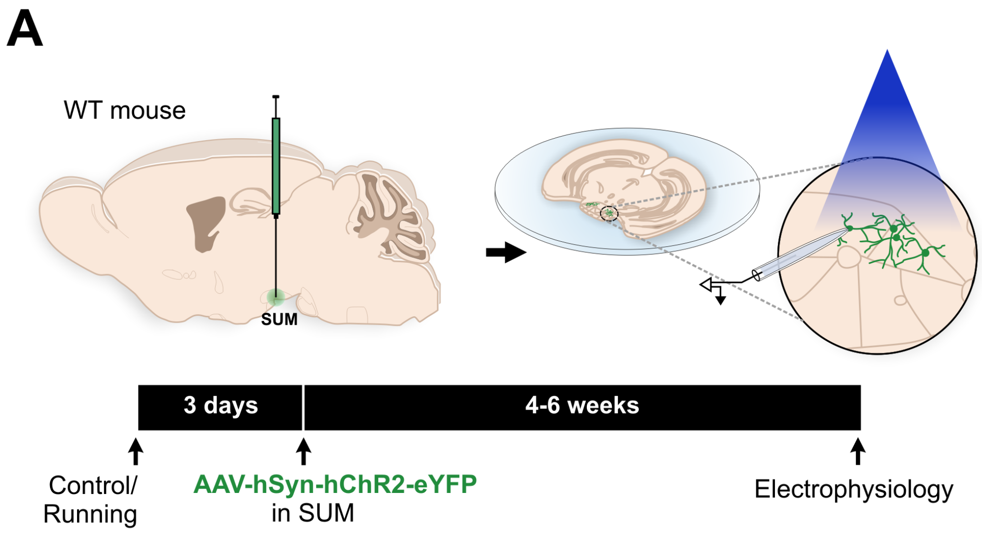

A
WT mouse
SUM
3 days
4-6 weeks
Control/
Running
AAV-hSyn-hChR2-eYFP
in SUM
Electrophysiology


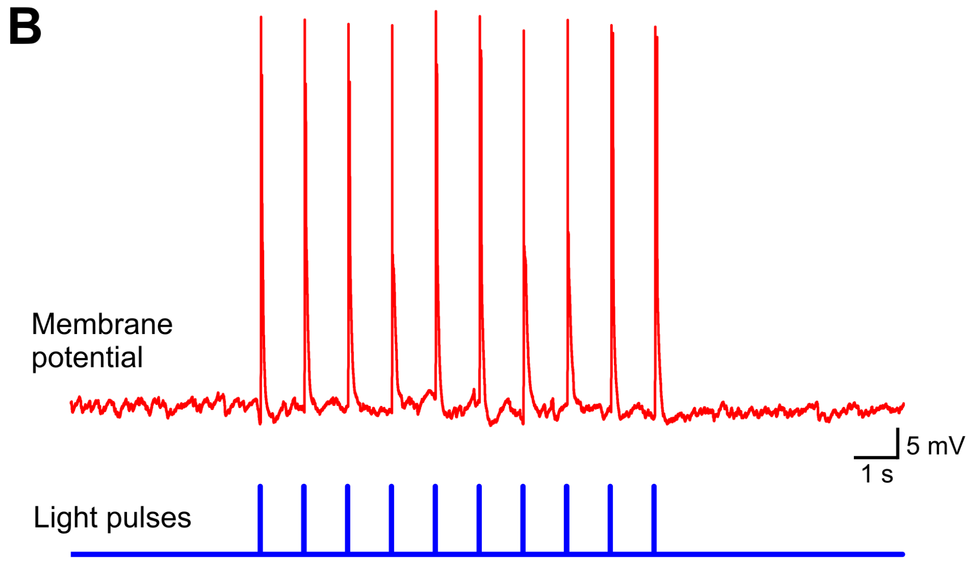

B
Membrane
potential
5 mV
1 s
Light pulses

**Fig. S3. Optogenetic stimulation elicited action potentials in SUM neurons expressing channelrhodopsin.** A) AAV2-hSyn-hChR2-eYFP was injected into the SUM of C57BL/6 mice. Patch-clamp recordings were performed after 4-6 weeks from ChR2-eYFP$^{+}$ SUM neurons. B) Whole-cell current-clamp recordings showing that blue light pulses (465 nm, 1 ms) elicit action potentials in ChR2-eYFP$^{+}$ SUM neurons.

### Supplementary Figure 4

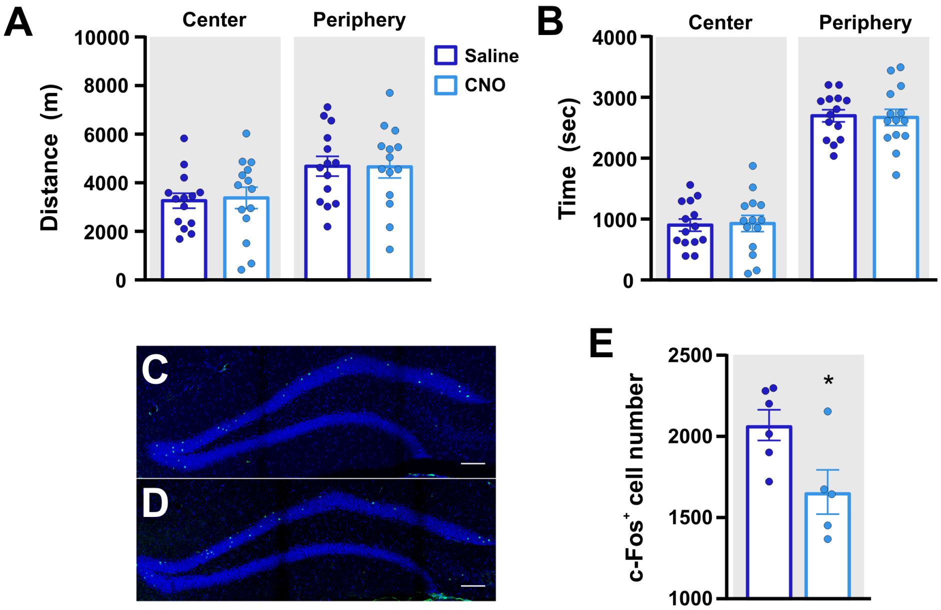


**Fig. S4. Open field behavior and c-Fos analysis**. (A, B) Open field activity. There was no significant difference across the groups in (A) distance travelled in the center ($t_{(13)}$ = 0.5066, $p$ = 0.6209) and periphery ($t_{(13)}$ = 0.1165, $p$ = 0.909) or (B) time spent in the center ($t_{(13)}$ = 0.3085, $p$ = 0.7626) or periphery ($t_{(13)}$ = 0.3086, $p$ = 0.7627) in the open field. (C, D) Photomicrographs of c-Fos$^+$ cells (green) in the dentate gyrus of the hippocampus in tissue derived from mice injected with (C) saline or (D) CNO. Nuclei are labeled with DAPI (blue). Scale bar = 100 µm. (E) Analysis shows a significant reduction in c-Fos$^+$ cells in the granule cell layer of the dorsal dentate gyrus ($t_{(9)}$ = 2.551, $p$ = 0.0311). Data are means ± SEM. Statistical analysis was performed using a paired *t*-test (A and B) and unpaired t-test (E).

#### Supporting Information

##### Experimental Procedures

##### Subjects

Male C57Bl/6J mice (Jackson Labs #000664) designated for electrophysiological experiments were individually housed at two months of age and randomly assigned to control or voluntary wheel-running conditions. Mice were housed in a 12h light–dark cycle (lights on at 6:00 a.m. and off at 6:00 p.m.) with food and water *ad libitum*. The experimental protocols were approved by the National Institute on Aging Institutional Animal Care and Use Committee (IACUC).

Adult male VGAT-ires-Cre mice (Jackson Labs #028862) were utilized for behavioral experiments. Mice were 13-15 weeks old (n = 35) and group-housed (3-5 per cage) in standard cages with a 12h light–dark cycle (lights on at 7:30 a.m. and off at 7:30 p.m.) with water and food *ad libitum*. Florida Atlantic University's IACUC approved all experimental procedures. All procedures were conducted in accordance with the guidelines required by the National Institutes of Health Guide for the Care and Use of Laboratory Animals.

##### Viral vectors

Retroviral vector RV-SYN-GTRgp, expressing nuclear green fluorescent protein (GFP), avian TVA receptor, and rabies virus glycoprotein (Rgp) driven by the neuron-specific synapsin promoter, retroviral vector CAG-GFP, retroviral vector CAG-RFP, and EnvA-pseudotyped rabies virus (EnvA-ΔG-MCh) were produced as previously described (*1-3*).

*Retrovirus.* To produce retrovirus, plates with 90% confluent human embryonic kidney (HEK293T) cells were transfected with vector (7.5 μg), CMV-GagPol (5 μg), and CMV-VSVG

(2.5 μg) using Lipofectamine 2000 (Invitrogen). Virus-containing supernatant was harvested 36 h later, filtered, and concentrated by ultracentrifugation. Virus titers were estimated to be ~ $1 \times 10^8$ i.u./ml by serial dilution into 293T cells.

*Rabies virus*. EnvA-pseudotyped Δgp-mCherry rabies virus (EnvA-ΔG-MCh) was produced by infecting the helper cell line, BHK-EnvARGCD, with glycoprotein-gene-deleted rabies virus vector (Δgp-mCherry) (Vector Core, Salk Institute). Supernatants containing rabies virus pseudotyped with envelope protein EnvA were harvested five days later, filtered, and concentrated by ultracentrifugation. Rabies virus titer was estimated to be ~ $1.2 \times 10^7$ i.u./ml and diluted for use to ~ $4 \times 10^6$ i.u./ml.

*AAV vectors*. AAV2-hSyn-hChR2-eYFP (titer: $5.7 \times 10^{12}$ vector genome (v.g.)/ml) and AAV5-CaMKII-ChR2-mCherry (titer: $5.7 \times 10^{12}$ vector genome (v.g.)/ml) were procured from the University of North Carolina vector core (Chapel Hill, NC, USA) and pAAV2-hSyn-DIO-hM4D(Gi)-mCherry ($5\times10^{12}$ vg/mL), from Addgene (#44362), aliquoted and stored at −80 °C until use.

**Stereotaxic surgery and viral vector injections**

After 3 days of housing in their respective conditions, mice underwent surgery during which they were anesthetized with Avertin (0.4 mg/g i.p.) and placed within a stereotactic frame (Stoelting Stereotaxic Instruments, USA). Retrovirus RV-SYN-GTRgp, CAG-GFP or CAG-RFP (1 μl) at a rate of 0.1 μl/min was injected into the right dorsal and ventral dentate gyrus (DG) using spatial coordinates relative to Bregma as follows: Dorsal DG, anterior–posterior (AP) = − 2.10 mm; medial–lateral (ML) = 1.9 mm; dorso-ventral (DV) = − 2.10 mm, and ventral DG, AP = − 3.10 mm; ML = 2.8 mm; DV = − 3.10 mm. Four to six weeks after RV-SYN-GTRgp injection, mice were anesthetized (Avertin 0.4 mg/g i.p.), and rabies virus EnvA-ΔG-MCh (1 μl) was delivered into the same DG locations. After 3-7 days, mice were sacrificed for further experiments. We injected 0.5-1 μl of AAV2-hSyn-hChR2-eYFP (for electrophysiology) or 0.3 μl of AAV5-CaMKII-ChR2-mCherry (for immuno-electron microscopy and co-localization studies) into the caudal hypothalamus (coordinates: AP = −2.7 mm, ML = 0, DV = −4.9 mm) and retroviral vector CAG-RFP (for electrophysiology) or CAG-GFP (for immuno-electron microscopy and co-localization studies) into the DG of the hippocampus (as described above). After a minimum of 4-6 weeks to allow for sufficient virus transport and infection, mice were used in electrophysiology experiments or perfused for histological analysis.

**Brain slice preparation and in vitro electrophysiological recordings**

Animals were deeply anesthetized with isoflurane and then rapidly decapitated. Brains were transferred to cooled (1–4 °C) dissection solution, containing (mM): 110 Choline-Cl, 2.5 KCl, 1.25 $NaH_2PO_4$, 25 $NaHCO_3$, 25 glucose, 1 $CaCl_2$, 7 $MgCl_2$, 0.6 $Na^+$ pyruvate, 1.3 $Na^+$ ascorbate, and 3 kynurenic acid. Horizontal hippocampal slices (300 μm thick) were obtained and transferred to a chamber containing incubation solution (in mM): 125 NaCl, 2.5 KCl, 1.25 $NaH_2PO_4$, 25 $NaHCO_3$, 2 $CaCl_2$, 2 $MgCl_2$, 20 glucose, 3 $Na^+$ pyruvate, and 1.3 $Na^+$ ascorbate in distilled water (equilibrated with 95% $O_2$ and 5% $CO_2$; pH 7.4). Slices were incubated at 34 °C for 10 min and stored at room temperature for >1 hour before patch-clamp experiments. Brain slices were submerged during electrophysiological recordings and superfused in artificial cerebrospinal fluid (ACSF) containing (in mM): 125 NaCl, 2.5 KCl, 1.25 $NaH_2PO_4$, 25 $NaHCO_3$, 2 $CaCl_2$, 1.3 $MgCl_2$, and 20 glucose (equilibrated with 95% $O_2$ and 5% $CO_2$; pH 7.4) at 30 ± 0.5 °C. Microelectrodes (5–8 MΩ) pulled from borosilicate glass were used for whole-cell recording and were filled with a K-based pipette solution containing (in mM): 130 K-gluconate, 7 KCl, 0.1 EGTA, 10 HEPES, 5 Mg-ATP, 0.5 Na-

GTP, and 10 Na-phosphocreatine (pH 7.4, adjusted with KOH). Biocytin (0.1%) was added to this internal solution for certain experiments.

Neurons in the DG were identified using infrared differential interference contrast video microscopy (Olympus BX51WI). Newborn neurons (4-6 weeks after retroviral labeling) were identified by RFP expression under epifluorescence. Mature granule cells born during development were recorded from the outer granule cell layer. Voltage-clamp and current-clamp recordings were performed using MultiClamp 700B amplifier system (Molecular Devices). Recordings were low-pass filtered at 2 kHz and digitized at 10–50 kHz (Digidata 1440 A and pClamp 10.4 Software; Molecular Devices). Data was acquired and analyzed using Clampfit 10.0 software (Molecular Devices) and Igor Pro (WaveMetrics Inc., USA). Series resistance was typically 10–30 MΩ.

Bicuculline methiodide (BIC, 100 µM; Abcam), picrotoxin (PTX, 100 µM; Abcam), (2S)-3-[[(1S)-1-(3,4-Dichlorophenyl)ethyl]amino-2-hydroxypropyl](phenylmethyl)phosphinic acid hydrochloride (CGP 55845, 1 µM), 6-Cyano-7-nitroquinoxaline-2,3-dione (CNQX, 10 µM; Abcam) and DL-2-Amino-5-phosphonopentanoic acid (AP5, 100 µM; Abcam) were added to the superfusion medium (ACSF) by dilution of their respective stock solutions.

**Electrophysiology data analysis**

All current clamp recordings were taken at the resting membrane potential (RMP) of the neuron. RMP was determined during the silent period of neuronal spontaneous activity. A current step protocol ranging from 20 pA to 240 pA of 600 ms duration was applied to determine the action potential firing profile and current injection relationship. Input resistance was determined from the slope of the linear regression fit through the V–I plot constructed by plotting the amplitude of the steady-state voltages (V) against the corresponding step current injections (I) from −10 to +5 pA of 1000 ms duration with 5 pA steps.

Membrane time constant was calculated by fitting a single exponential to the initial 100 ms of membrane potential deflection. Capacitance was calculated as the ratio of the membrane time constant and membrane resistance. Depolarizing current ramp ranging from −60 to +150 pA, for an interval of 1.1 s, was used in current clamp mode to elicit action potentials. The threshold was analyzed from the first action potential. Action potential amplitude and half-width were calculated from the first action potential generated by a step pulse of 600 ms with minimal current injection of +20 pA. Action potential amplitude was defined as the difference between the membrane potential at the inflection point to the peak of the action potential. Spike frequency was calculated as the reciprocal of the mean of the inter-spike intervals generated during 0.6 s long step depolarizations.

Synaptic events were examined using Igor Pro (Wave Metrics Inc., USA). Event detection was performed using amplitude threshold crossing that were validated after visual inspection. For spontaneous synaptic current, >200 s of raw trace was evaluated from each neuron recorded in voltage clamp mode at a holding potential of -50 mV, where the inward and outward synaptic currents represented excitatory postsynaptic currents (EPSCs) and inhibitory postsynaptic currents (IPSCs), respectively. All the EPSCs and IPSCs were analyzed separately for peak amplitude, frequency, rise time, and decay time constant in each neuron, which were averaged for each neuron and used for further statistical analysis. A single exponential function was fitted to decay times of individual events to evaluate the decay time constant. Peak amplitude of spontaneous synaptic events was determined relative to the baseline current value. The 20–80% rise-time was calculated between the start and the peak of a synaptic event.

#### Optically evoked synaptic stimulation

For optical stimulation, light was produced by a 465-nm Plexibright LD-1 LED module (Plexon, Dallas, TX, USA) and delivered by an optical fiber (200 µm core diameter, NA = 0.5) that was submerged in ACSF and placed ~300 µm from the recording site. SUM axons expressing ChR2 were activated with the optical fiber placed just above the DG granule cell layer. Square pulse stimulation (1 ms, 20 mW) was applied at 0.1 Hz using digital commands from the Digidata 1440A (Molecular Devices) and PlexBright LD-1 Single Channel LED Driver (Plexon, Dallas, TX, USA). Optical stimulation in DG evoked synaptic currents that were measured in voltage clamp mode at different holding potentials: -90, -80, -70, -60, -50 and -40 mV. Optically evoked synaptic membrane potentials were also measured in current-clamp mode at resting membrane potential and with depolarizing current injection (20-180 pA in steps of 20 pA for 1340 ms) to measure subthreshold and suprathreshold effects. Depolarizing step current injections elicited spiking activity that was recorded. Action potential counts 50 ms before and after optical stimulation were compared to examine the immediate effect of brief (1 ms) SUM fiber stimulation.

#### Electron microscopy and immunohistochemistry

Methods were described previously (*4*). Briefly, vibratome tissue sections from three male WT-ChR2-mCherry-GFP mice (C57BL/6 mice injected with AAV5-CaMKII-ChR2-mCherry in SUM and CAG-GFP in DG, respectively; Fig. 2A) were rinsed and incubated with 1% sodium borohydride to inactivate free aldehyde groups, rinsed, and then incubated with blocking solution. Sections were then incubated with primary antibodies [mouse anti-mCherry (1:1,000, Takara, #632543), guinea pig anti-VGAT (1:500, Frontier Institute, VGAT-GP-Af1000, RRID: AB_2571624) and rabbit anti-GFP (1:2,000, Frontier Institute, GFP-Rb-Af2020, RRID:

AB_2571573)]. All primary antibodies were diluted with 1% normal goat serum (NGS), 4% BSA in PB supplemented with 0.02% saponin, and incubations were for 24 h at 4°C. Sections were rinsed and incubated overnight at 4°C in the corresponding secondary antibodies. Sections were rinsed in PB, and then in double-distilled water, followed by silver enhancement of the gold particles with the Nanoprobe Silver Kit (2012, Nanoprobes) for 7 min at room temperature. Next, sections were incubated in avidin-biotinylated horseradish peroxidase complex in PB for 2 h at room temperature and washed. Peroxidase activity was detected with 0.025% 3,3'-diaminobenzidine (DAB) and 0.003% H2O2 in PB for 5-10 min. Sections were rinsed with PB and fixed with 0.5% osmium tetroxide in PB for 25 min, washed in PB followed by double distilled water and then contrasted in freshly prepared 1% uranyl acetate for 35 min. Sections were dehydrated through a series of graded alcohols and with propylene oxide. Afterwards, they were flat embedded in Durcupan ACM epoxy resin (14040, Electron Microscopy Sciences). Resin-embedded sections were polymerized at 60 °C for 2 days. Sections of 60 nm were cut from the outer surface of the tissue with an ultramicrotome UC7 (Leica Microsystems) using a diamond knife (Diatome). The sections were collected on formvar-coated single slot grids and counterstained with Reynolds lead citrate. Sections were examined and photographed using a Tecnai G2 12 transmission electron microscope (Fei Company) equipped with the OneView digital micrograph camera (Gatan).

**Ultrastructural analysis of brain tissue**

Serial ultrathin sections of the DG from 3 male WT-ChR2-mCherry-GFP mice. Synaptic contacts were classified according to their morphology and immunolabeled and photographed at a magnification of 6,800-13,000×. The morphological criteria used for identification and classification of cellular components or type of synapse observed in these thin sections were as previously described (*4*). In the serial sections, a terminal containing greater than 5 immunogold particles were considered as an immuno-positive terminal. Pictures were adjusted to match contrast and brightness by using Adobe Photoshop (Adobe Systems Incorporated, Seattle, WA). This experiment was successfully repeated three times.

**Fluorescence microscopy and image analysis.**

Methods were described in a previous study (*5*). Briefly, free-floating coronal sections (40 µm) from 3 male WT-ChR2-mCherry-GFP mice (n = 3) were incubated for 1 h in PB supplemented with 4% BSA and 0.3% Triton X-100. Sections were then incubated with cocktails of primary antibodies: mouse anti-mCherry (1:500) + guinea pig anti-VGluT2 (1:500) + rabbit anti-VGAT (1:500, Frontier Institute, VGAT-Rb-Af500, RRID: AB_2571622) + goat anti-GFP (1:2,000, Frontier Institute, GFP-Go-Af1480, RRID: AB_2571574) overnight at 4ºC. After rinsing 3 × 10 min in PB, sections were incubated in a cocktail of the corresponding fluorescence secondary antibodies (1:100): Alexa-Fluor-405-anti-guinea pig + Alexa-Fluor-488-anti-goat, Alexa-Fluor-594-anti-mouse + Alexa-Fluor-647-anti-rabbit for 2 h at room temperature. After rinsing, sections were mounted on slides. Fluorescent images were collected with an Olympus FV1000 Confocal System (Olympus). Images were taken sequentially with different lasers with 10× and 100× oil immersion objectives and Z-axis stacks were collected at 0.2 µm. Amira microscopy software (Thermo Fisher Scientific) was used to analyze z-stacks of confocal images from 3 WT-ChR2-mCherry-GFP mice (64 × 64 × 10 µm for each image; four images from each animal) to obtain 3D reconstructions of putative synapses. This experiment was successfully repeated three times.

**Behavioral assays, General Procedure / Overview**

Adult male VGat-ires-cre mice (Jackson Labs #028862), 13-15 weeks old at the beginning of the experiment (n = 35) were group-housed (3-5 per cage) in standard cages with *ad libitum* access to food and water on a standard 12h/12h day/night cycle: lights were switched on at 7:30 AM and off at 7:30 PM. Mice (n = 35) underwent stereotaxic surgery to inject pAAV2-hSyn-DIO-hM4D(Gi)-mCherry into the SUM nucleus, resulting in the attrition of 5 mice. Four weeks thereafter, mice (n = 30) started habituation and pre-training in the touchscreen task. Mice were food-restricted from 8 days to ~4 grams of chow per day, starting before the beginning of the touchscreen experiment. Exclusion criteria resulted in n = 22 mice for task training, which was successfully completed by n = 20 mice. These mice were divided into saline control and CNO groups (n = 10 per condition), for two pattern separation probe trial sessions of 16 days each. Mice who received CNO in the first probe trial subsequently received saline, and mice who received saline subsequently received CNO for the second set of probe trials. Following completion of each set of probe trials, mice were evaluated in the open field test following saline or CNO injection. Next, mice were trained in the Morris water maze for 9 days and then tested for memory retention in a probe trial 24 h after the last training trial following saline or CNO injection. Finally, mice were injected with saline or CNO and tested in the contextual fear conditioning paradigm (Figure 4A). Following completion of behavioral testing and before brain tissue collection, mice were injected with saline or CNO and housed in a new cage with a running wheel for one hour for c-Fos activation. Immediately thereafter, mice were deeply anesthetized and perfused transcardially. Histological analysis of the injection site location and spread of the vector resulted in the inclusion of n = 14 mice in the final data analysis.

**Surgery**

Mice 13-15 weeks old, underwent stereotaxic surgery. Mice were anesthetized with isoflurane (Abbott) and stereotaxic surgery was performed to deliver 0.2 □l of pAAV2-hSyn-DIO-hM4D(Gi)-mCherry (≥ $5\times10^{12}$ vg/mL, Addgene, #44362-AAV2) into the supramammillary nucleus using spatial coordinates relative to Bregma as follows: AP = -2.70 mm; ML = 0.00 mm; DV = -4.90 mm.

**Drug Administration**

Clozapine N-oxide (CNO) (freebase) (HelloBio, HB1807) was dissolved in 0.9% saline at 1mg/ml and administered by intraperitoneal (i.p.) injection at 5 mg/kg. CNO or an equivalent volume of 0.9 % saline was administered daily, 30 min prior to the start of probe trial testing for the pattern separation task in the touchscreen, and similarly prior to open field behavior, water maze training and testing, and in the fear conditioning paradigm before habituation, conditioning, tone-cued and context testing.

**Touchscreen-based pattern separation test**

The testing apparatus for the location discrimination (LD) task (Lafayette, Cambridge) includes a sound attenuating cubicle (width 56.5 cm × depth 53.2 cm × height 54.5 cm) and a touch screen chamber (width at feeder 4.6 cm × width at screen 23.8 cm × depth 17 cm × height 23 cm) consisting of a black Plexiglas trapezoidal wall, touch screen installed with infrared sensor (WhiskerServer Controller), a perforated-metal floor, reward area, a liquid pump feeder, a speaker with tone generator, a tray light and a house light. A black Plexiglas "mask" panel, which includes two rows of six square windows (2.5 cm × 2.5 cm, equally spaced 1 cm apart), located 1.6 cm from the bottom of mask panel was inserted in front of the touch screen to allow for presentation of white stimuli on the 6 windows at the bottom of the mask panel. When an animal touches the white stimulus coming from one of the 6 windows, the software (ABET II Touch software) counted and saved its activity automatically.

*Pre-training (7 weeks and 3 days):* Mice were given *ad libitum* access to water, but food-restricted from 8 days before the beginning of the LD task throughout the touchscreen experiment. Mice were food-restricted to ~4 gr of chow per day. Mouse body weights were taken daily. Mouse weight was 22 (+/- 0.87) grams at the onset and 24 (+/- 1.14) grams at completion of the experiments. Habituation: For the first 2 days of shaping, mice were placed in the touchscreen chambers for 20 min each day to become familiar to the chamber surroundings. In the chamber the light was turned on, the screen was off, and the reward trough contained condensed milk diluted 25:75 with distilled water (Sweetened Condensed Milk, Eagle Brand). Pavlovian training (Initial touch training): On day 3, mice continued onto Pavlovian initial touch training. The stimulus was a white square that was displayed randomly in one of the windows on the bottom row while the remaining windows were blank. The white square was not shown in the same window more than three times in a row. When the mouse touched the white square, a drop of reward and a tone were provided with illumination of the food tray light. Once the mouse collected the reward, the tray light turned off and triggered the inter-trial interval (ITI, 10 s) for the next trial. The length of one trial was from the onset of the stimulus to the end of the ITI. Following the ITI, the next white square was displayed in one of the bottom row windows. Mice were trained until they reached a criterion of 30 trials in 1 h. "Must touch" stage: A trial started with the stimulus presented randomly in one of the bottom row windows, while the other windows remained blank. Upon touching the stimulus, the tray light, the tone, and a drop of reward were delivered, followed by a 10 s ITI. After the ITI, the next trial started. A touch to a blank window was considered as "incorrect" and no reward or tone were provided. Mice were trained until they reached a criterion of 30 trials in 1 h. “Must initiate” stage: The stimulus was presented in the same way as in the “Must touch” stage and a touch to the stimulus resulted in the tone and reward response with the tray light. Following a 10 s ITI after obtaining the reward, mice were required to poke their nose into the reward trough again to start the next trial. Once a mouse had reached a criterion of 30 trials in 1 h, the next stage started. “Punish Incorrect” stage: The stimulus and reward were given in the same way as described above. However, if a mouse touched the blank window, instead of the white window, the house light in the ceiling of the chamber was inverted for a time-out period of 5 s and no reward was supplied. After the time-out period was over, the house light was reverted, and the 30 s ITI period started. Mice were trained until a criterion of 23 correct trials out of 30 total trials in 30 min (85% correct trials) for two consecutive days was reached.

*Task training (19 days)*: At the beginning of each trial, two white square stimuli were displayed at positions 2 and 5 out of the six windows, with position 1 being the left-most window. For each trial, one of two stimuli were supported by both reward and tone. The side of the correct stimulus was randomly counterbalanced between mice. When a mouse reached the performance criterion (seven trials out of eight trials correct) the reward-support stimulus was reversed, in which the previous incorrect window became the support stimulus and vice versa. This training lasted for 60 trials or until eight reversals were reached, whichever was first. The next day, animals were trained the same way and the originally correct window was the same one as in the last session of the day before. Mice were trained until they reached the acquisition performance criterion, 7 out of 8 trials correct, for 3 out of 4 days in a row.

*Probe trials (2 x 16 days)*: For the probe trial sessions, mice were injected with either saline or CNO (5 mg/kg) at 30 min prior to the onset of testing. The probe consisted of the presentation of either a small or a large separation between stimuli, which was counterbalanced between mice. In the large separation trial, two stimuli were displayed at the same time in windows 1 and 6, the most left and right side of 6 windows respectively, and the two stimuli were displayed in window 3 and 4 in the small separation trial. For 60 min, unlimited trials were given. Each separation was interchanged every 2 days over 16 days of testing. For example, if a mouse began with the large separation, this continued for 2 days, first and second days, and then the small separation was presented on the third and fourth days. This pattern was repeated for 16 days of probe trial testing. A second 16-day probe trial was performed with the same group of mice, switching mice that received CNO to saline, and mice that received saline to CNO, respectively. The number of trials to criterion (7 out of 8 trials correct) and the number of reversals were used for the data analysis.

**Open field test**

Spontaneous locomotor activity was evaluated after the first and second rounds of 16-day probe trials by using an infrared photobeam-controlled open-field testing chamber (Version 4.0, Med Associates, St. Albans, VT, USA; height 20.3 cm, width 27.3 cm, depth 27.3 cm). The center zone was defined as an area of 14.29 cm × 14.29 cm. Mice were injected with either saline or CNO (5 mg/kg) 30 min before they were placed in the open field chamber and explored for 60 min. Mouse movement was detected by three 16-beam infrared arrays installed on X, Y, and Z axes and the time and distance in the center and periphery of the open field were automatically recorded by Activity Monitor software.

**Morris water maze**

Spatial learning and retrieval were evaluated using the Morris water maze paradigm (*6*). The apparatus consisted of a pool filled with water (24-26 °C), made opaque with white nontoxic paint

(Tempura, Crayola), and contained a platform that was hidden 1 cm below the surface of the water. The pool was surrounded by a black curtain, 60 cm from the edge of the pool to which visual cues were attached. Indirect illumination of both the pool and cues (65 lux at the center of the maze) was achieved by positioning 4 white lights equidistantly around the pool and facing the ceiling. The pool was virtually separated into 4 quadrants, called northeast (NE), northwest (NW), southeast (SE) and southwest (SW). Mice were acclimated to the testing room daily for at least 30 min before starting the test. Mice were subjected to one acquisition session per day for 9 days. Each session consisted of four 60 s trials with a 15 s inter-trial interval, in which the hidden platform was in the NE quadrant of the pool. Mice were placed at a different starting point for each trial. If the mouse found the platform, the trial was immediately terminated. Mice were placed on the platform if they did not find it within the allocated time. In either case, mice were allowed to explore the visual cues for 15 s while on the platform. Spatial memory retention was evaluated in a 60 s probe trial (24 h after completion of the last acquisition session) in which the platform was removed. For the probe trial, mice started in the SW quadrant. Data was collected by an automated video tracking system (Ethovision XT 17.5).

**Fear Conditioning**

Pavlovian conditioning-based approach was used to evaluate fear memory. Mice were tested within a four-chamber system utilizing a near-infrared video conditioning system (MED Associates Inc., Fairfax, VT, USA) that allows for testing of four animals simultaneously. Each chamber consists of stainless-steel walls with one transparent plexiglass door and a floor of parallel stainless-steel rods connected to a shock generator. The chamber has an overhead white light (SKU ENV-229M) to illuminate the inside (3 to 8 lux), and a speaker which is used to deliver a tone, to serve as the conditioned stimulus (CS). Each of these chambers was located within a larger noise-attenuating chamber (height 31.75 cm x width 71.12 cm x depth 59.69 cm), including a ventilation fan delivering background noise. A near-infrared camera-tracking system (MED Associates, Georgia, VT, USA) was used to automate measurement of freezing behavior (Motion Threshold (au) = 20; Detection Method = Linear; Min Freeze Duration (f) = 18). The paradigm consisted of four phases: (i) Habituation, (ii) Conditioning, (iii) Tone-cued, and (iv) Contextual phase. Mice were acclimated to the testing room for at least 1 h before test onset. Habituation (Day 1) consisted of a single 330 s session in which mice were placed within the chamber and allowed to freely explore the environment. A 1% liquinox solution was in the tray located underneath the stainless-steel rods to serve as an odor cue and to facilitate contextual perception and discrimination. Freezing behavior was measured for the entire 330 s. Mice were immediately returned to their home cages at the end of the session. The inner surfaces of the chamber and the tray were cleaned thoroughly with 70% isopropyl alcohol. Conditioning phase (Day 2) was similar to the Habituation session with the exception that a CS, tone, paired with an Unconditioned Stimulus (US), foot shock, was presented 60 s after the beginning of the trial. CS consisted of a tone (5000 Hz, 90 dB tone) for 30 s, that ended with the presentation of the US, a 0.5 s foot shock (0.5 mA) delivered 0.5 s before the end of the CS (TS1). Thereafter, the CS-US was repeated two more times (TS2 and TS3, respectively) with a 90 s inter-stimulus interval. Freezing behavior was measured during the same 30 s in which the tone was on. CFR, the response evoked by the CS-US association, was considered to be formed if freezing behavior was higher during the third than the first CS-US presentation. Tone-cued phase (Day 3) was performed to evaluate long-term evocation of Pavlovian conditioning. It consisted of a single session similar to the Conditioning phase, except that no US was delivered. In addition, to avoid contextual (visual, tactile and olfactory) influence on the CS-US association, the original environment was modified by adding a blue semitransparent plastic film on top of the plexiglass chamber, an arch-shaped white plastic panel within the chamber and a white plastic panel at the base of the chamber as well as the replacement of the 1% liquinox solution by an 1% acetic acid solution. Two Contextual sessions (Day 3 - one hour after Tone-cued phase – and Day 5) were performed to evaluate long-term evocation of contextual CFR (i.e., whether freezing behavior was higher during the tests than during Day 1). These were identical to the Habituation session.

**Immediate early gene expression**

To validate the use of CNO we quantified c-Fos in the DG granule cell layer. After completion of behavioral testing, animals were given an injection of either saline or CNO (5 mg/kg). Thirty min after the injection, mice were individually housed in a new cage with access to a running wheel for one hour. Immediately thereafter mice were deeply anesthetized and perfused transcardially.

**Perfusion/Histology**

Mice were injected with an overdose of anesthetic drugs (Ketamine 90-120 mg/kg, xylazine 7.5-10 mg/kg) and perfused transcardially with 0.9% saline at room temperature (followed by cold (4 $^{o}$C) 4% paraformaldehyde (PFA)). After perfusion, brains were dissected, post-fixed in PFA for 24 h, and afterwards equilibrated in 30% sucrose. Consecutive coronal sections (40 μm) were taken using a freezing microtome (HM450, ThermoFisher Scientific, Waltham, MA, USA) through the rostral-caudal extent of the brain and stored in phosphate buffered glycerol at −20 ◦C as described (*7*).

**Immunohistochemistry**

A one-in-six series of free-floating sections (40 μm) was washed in tris-buffered saline (TBS). Following washes in TBS sections were blocked with $TBS^{++}$ (3% Donkey serum in TBS, triton ($TBS^{++}$) and then incubated for 72 h in primary antibodies, (RFP rabbit polyclonal, 1:250, Rockland; c-Fos mouse monoclonal, 1:250, Abcam). After a 72 h incubation, tissue was washed in TBS, followed by $TBS^{++}$ and then incubated in the secondary antibody solution (donkey-anti-rabbit CY3; donkey-anti-mouse AF488; 1:250, JacksonImmunoResearch). Thereafter, the tissue was stained for DAPI (1:10000), mounted and coverslipped with DABCO-PVA. DG c-Fos positive cells were counted in a one in six series of sections using a fluorescence microscope and 20x objective (Olympus, BX51).

**Statistical analysis**

Statistical analyses were performed using GraphPad Prism 7 (GraphPad Software Inc, CA). ANOVA was used for multiple comparisons and the Tukey's test or Fisher's PLSD test for the post-hoc analysis. Student's t-test was conducted for normally distributed data. Two-sample Kolmogorov–Smirnov test was conducted to compare population distributions. Averaged data are shown as mean ± SEM and statistical significance was concluded at $p < 0.05$ for all analysis.
